# Development of an undergraduate cell biology laboratory to assess pigmentation and cell size in a zebrafish model of uveal melanoma

**DOI:** 10.1101/2023.09.15.558009

**Authors:** Andrea M. Henle

## Abstract

This study outlines a two-week laboratory module for an authentic cell biology undergraduate research experience that uses zebrafish (*Danio rerio*), a popular model organism for research. Previous research has indicated that course-based undergraduate research experiences such as this one increase student confidence, active learning, and retention. During this research experience, students investigate variations in pigmentation in the caudal fins of wild type and transgenic fish (*Tg(mitfa:GNAQ*^*Q209L*^)). The transgenic fish express a hyperactive Gα protein, GNAQ^Q209L^, under the melanocyte-specific mitfa promoter, offering insights into uveal melanoma, a common eye cancer. Students specifically analyze the black pigmented cells, melanophores, within the caudal fin. We determined that the transgenic zebrafish have increased pigmentation in their caudal fins but smaller melanophores. These results suggest there are more melanophores in the *Tg(mitfa:GNAQ*^*Q209L*^) fish compared to the wild type. Future undergraduate research could investigate these cellular differences. This research experience imparts microscopy and image analysis skills and instills the ability to grapple with large datasets, statistical tests, and data interpretation in alignment with biology education principles. Post-lab surveys reveal students attain confidence in the above skills and in handling animals, along with a deeper appreciation for model organism research and its relevance to cancer cell biology.

## Introduction

The zebrafish (*Danio rerio*) is a popular model organism not only in scientific research laboratories^1–4^, but also in undergraduate education settings^5,6^ and for K-12 outreach^7–11^. This small freshwater fish is used as a model across many subfields of biology and biomedical science, including cell biology, developmental biology, genetics, neuroscience, immunology, and cancer biology^12^. Zebrafish have been adopted by laboratories and schools worldwide because of their fast development, translucency during embryonic and larval stages, ease of genetic manipulation, and affordability to maintain in large numbers^13^.

Students in the former sophomore-level course, Cell and Molecular Biology, and the current upper-level course, Advanced Cell Biology, at Carthage College (Kenosha, WI) use zebrafish to conduct authentic cancer and cell biology research during a two-week lab module. The laboratory portion of the course meets for 3 hours, once per week. This laboratory research experience introduces students to using a model organism for cell biology research, provides them with microscopy skills, and introduces them to analysis of microscopic images through the freely available image processing software, FIJI (fiji.sc). Another major aspect of this laboratory involves grappling with the ambiguity of a large dataset. The data from all of the students is compiled into one class dataset, and students must perform statistical tests to identify outliers, calculate means, standard deviations, and p-values. Lastly, students use their analyzed data to generate professional scientific figures and communicate their findings. These learning objectives are in line with the core concepts and competencies put forth by AAAS in their document, “Vision and Change in Undergraduate Biology Education: A Call to Action”.^14^

Students meet the learning objectives of the lab by investigating differences in the area of pigmentation in zebrafish caudal fins and in the size (µm^2^) of individual pigmented cells within the caudal fin. Specifically, students in this laboratory investigate the black pigmented cells called melanophores. Each student or pair of students analyzes pigmentation in at least two different zebrafish backgrounds: wild type and transgenic (*Tg(mitfa:GNAQ*^*Q209L*^). The transgenic fish used at Carthage College have a copy of human GNAQ^Q209L^ (a Gα protein) under the control of the melanocyte specific promoter, mitfa. Much of this laboratory experience at Carthage College discusses the role of the GNAQ protein in uveal melanoma and demonstrates how the zebrafish model organism is used for cancer research.

Uveal melanoma is the most common eye cancer in humans with an estimated 1700 cases diagnosed yearly in the United States^15,16^. Patients diagnosed with this disease typically undergo surgery or radiation for treatment. However, uveal melanoma is an aggressive cancer and the majority of patients develop untreatable metastatic disease in the liver. There is currently no treatment for this recurrent form of the disease. Activating point mutations in two genes, GNAQ and GNA11, are implicated in over 83% of patients with this disease^17^. However, the exact role of these two mutated and oncogenic Gα proteins needs further elucidation.

The zebrafish model organism is used to study uveal melanoma because zebrafish have similar genomes to humans, tumors develop quickly (i.e. within months when a driver mutation such as p53-/-is also present (*tp53*^*M214K/M214K*^))^18^, and they have similar eye anatomy. Through this laboratory research experience, students learn about the role of mutations in genes and how a hyperactive Gα subunit in a G protein interferes with downstream signaling pathways. In particular, the students investigate how the hyperactive Gα subunit alters cell size and overall pigmentation in the caudal fin of the zebrafish.

Of importance, this laboratory does not need to focus on cancer and lends itself quite easily to the analysis of cellular changes in other zebrafish pigmentation backgrounds, including the commonly available *leopard*^19^, *jaguar*^20^, *panther*^21^, and *golden*^22^ zebrafish backgrounds.

## Methods

### Fish husbandry

Wild type *(TAB5/14)*^23^ and transgenic *Tg(mitfa:GNAQ*^*Q209L*^)^24^ zebrafish were maintained according to standard operating procedures and guidelines^25,26^. All zebrafish protocols were approved by the Institutional Animal Care and Use Committee at Carthage College. Each student completed CITI training prior to working with zebrafish and completed an Occupational and Health Safety form to gain approval for conducting research with zebrafish.

### Anesthetization of zebrafish and caudal fin amputation

Adult zebrafish were individually anesthetized in 0.05% tricaine (MS-222) 0.0625% sodium bicarbonate, pH 7. Upon anesthetization, zebrafish were placed directly on a glass slide and their caudal fin was amputated using a clean scalpel. Each fish was returned to a clean tank with fresh aquarium water to recover from the surgery. The scalpel was used to spread out the caudal fin on the microscope slide. Excess solution was blotted off the slide with a kimwipe. Cover slips were not placed on the slides as it was discovered that the bony rays in the caudal fin interfered with the imaging and depth of field when a cover slip was placed on top of the fin.

### Microscopy to image melanophores

Amputated caudal fins were immediately imaged after surgery on a Nikon SMZ18 stereomicroscope. Magnifications ranging from 0.75X to 13.5X were used to capture at least one image of the entire caudal fin and at least one image showing melanophores within three different possible regions (dorsal, medial, or ventral) of the caudal fin at high magnification. Scale bars were added in NIS Elements software and images were captured with a DS-Ri2 color CMOS camera.

### Quantitative Image Analysis

Students used FIJI (Fiji is Just Image J) freeware to perform two different types of image analysis. The scale bar for each image was set in FIJI and images were converted to 8-bit grayscale format. For the first analysis, students used the freehand selection tool to outline the entire caudal fin, and the measure and thresholding tools to calculate both the area of black pigmentation within the entire caudal fin and the overall area of the entire caudal fin. The percent of pigmentation within the overall fin was then calculated in Microsoft Excel.

For the second image analysis, students used FIJI to approximate the size of at least 10-15 individual melanophores within their magnified image of the caudal fin. Students used the freehand selection tool to separately circle individual melanophores within ImageJ and again used the measure and threshold tools to calculate the area of black pigmentation (melanin) within the selection. These pigment dispersion measurements are a rough approximation of individual cell size since we did not use a membrane marker for the melanophores. These measurements assume that the melanin pigment is dispersed throughout the entire cell, which may not always be the case.

### Statistical Analysis and Data Presentation

Each student’s image analysis data was compiled into a class data spreadsheet in Microsoft Excel for the second week of the laboratory. Students used a Grubb’s test (free outlier calculator) from GraphPad (https://www.graphpad.com/quickcalcs/Grubbs1.cfm) to help them sort the data and remove outliers.

Students were also encouraged to speak with each other during the laboratory to address potential outliers and to ask each other if a technical part of the analysis may have been performed incorrectly. Students then calculated the sample size, mean, standard deviation, and standard error of the mean for each dataset in Excel.

Students performed an unpaired t-test with the quick calculator from GraphPad (https://www.graphpad.com/quickcalcs/ttest1/?format=SD) to determine any difference between the mean pigmented area for each background of fish or the average area of pigment dispersion (an approximation of cell size) within the different backgrounds of fish. A p-value <0.05 was deemed statistically significant.

In the laboratory, students used Microsoft Excel to generate bar graphs of analyzed data. All graphs in this manuscript were prepared with Prism 7.

### Post-Lab Survey

Students were asked to complete an anonymous post-lab survey after completion of the second week of the laboratory. Students were informed their participation in the survey was completely voluntarily and would not affect their grade. This study was deemed exempt by the Carthage College Institutional Review Board.

### Instructor Notes

Instructors should be prepared to discuss several topics with the students at the start of lab and throughout this laboratory investigation. First, instructors should review G-protein coupled receptors and their downstream activation. This leads nicely into the discussion of the GNAQ gene which codes for the alpha subunit of a G protein. Instructors can discuss the role of the point mutation in the hyperactivation of the Gα protein and its inability to hydrolyze GTP to GDP. This can provide an opportunity to compare oncogenes and tumor suppressors, if desired.

Subsequently, instructors should be prepared to lead the class on the generation of hypotheses. If the Gα protein is always active in melanophores in zebrafish or uveal melanocytes in humans, what effect might that have on these cells? Students typically propose that the cells will proliferate or divide more frequently, have a higher survival rate, grow in size, or migrate further distances away from or within the tissue of origin.

Instructors can provide some background on the zebrafish model organism, its advantages, and limitations. They can discuss what a transgene is, how a transgenic model is made in zebrafish (e.g. through injection of DNA constructs into single cell embryos followed by screening and several rounds of mating the progeny). Instructors should also specifically address that the genotype of zebrafish used in this laboratory expresses the human GNAQ^Q209L^ gene under the control of the melanocyte-specific mitfa promoter. It is also helpful at this point to provide students with information about the three types of pigmented cells in zebrafish: iridophores (iridescent), xanthophores (yellow pigmented cells), and melanophores (black pigmented cells).

Instructors can then ask students to work in small groups to design an experiment that will address one of the previously stated hypotheses and will utilize the zebrafish model organism. If following along with this manuscript, the instructor should attempt to get the class to agree to doing an experiment that investigates the differences between pigmented area in Tg vs WT zebrafish and measuring the size of the melanophores (or the area of pigment dispersion within each individual melanophore) in Tg vs WT fish. These experiments will allow students to observe the direct effect of expression of GNAQ^Q209L^ in the melanophores. Specifically, the latter experiment allows them to address whether the GNAQ^Q209L^ transgene causes cells to grow in size.

## Results and Discussion

GNAQ is a known oncogene associated with uveal melanoma. Therefore, we asked whether its expression in melanophores results in changes in pigmentation in adult zebrafish. As expected, adult *Tg(mitfa:GNAQ*^*Q209L*^) fish (Tg) display significantly increased pigmentation in their caudal fins compared to their wild type (WT) clutchmate controls (Figure 1). The percent of pigmentation across the entire caudal fin is increased (Figure 2). These findings corroborate a previous study^24^ which found increased pigmentation in these transgenic fish at the embryonic stage.

**Figure 1.**
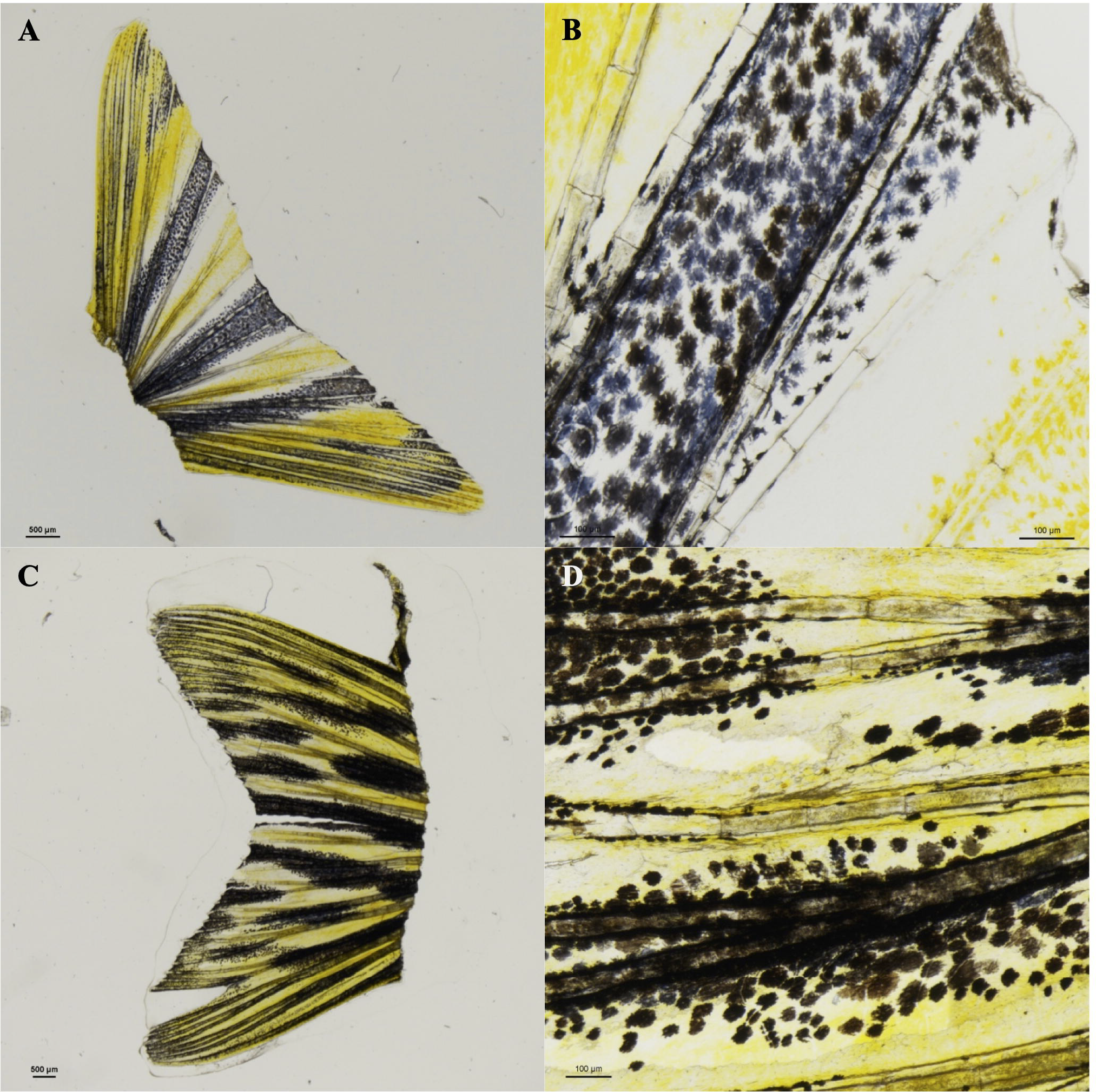
Pigmented cells (melanophores) in the caudal fins of adult wild type (*WT*) vs transgenic (*Tg(mitfa:GNAQ*^*Q209L*^)) (Tg) zebrafish. A.) amputated WT caudal fin. Scale bar = 500 µm. B.) WT caudal fin melanophores. Scale bar = 100 µm. C.) amputated Tg caudal fin. Scale bar = 500 µm. D.) Tg caudal fin melanophores. Scale bar = 100 µm. Images were captured on a Nikon SMZ18 stereoscope.

**Figure 2.**
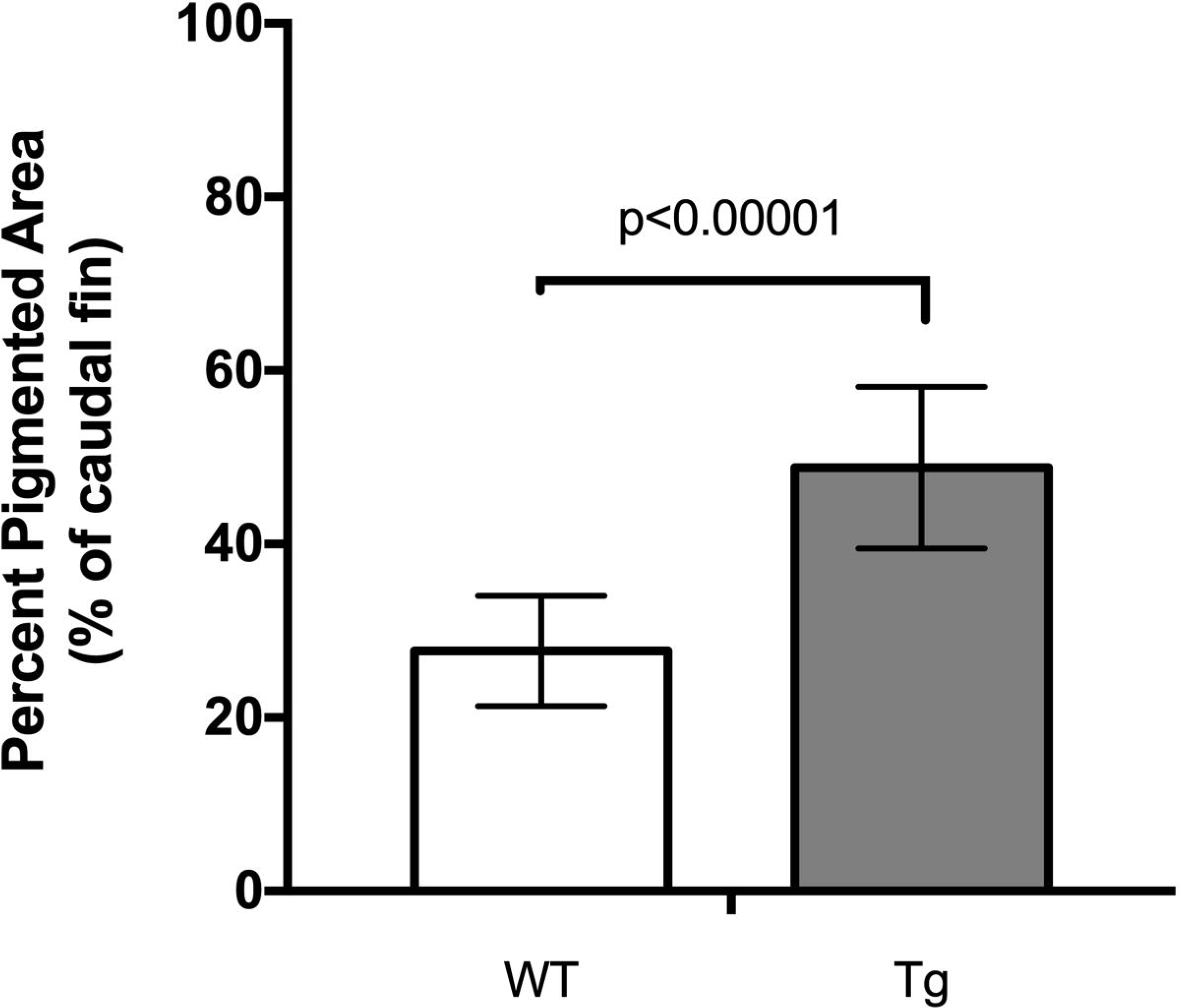
Average percent pigmentation in the caudal fins of adult WT and Tg zebrafish. Images of the caudal fins of adult zebrafish were captured on a Nikon SMZ18 stereoscope. Image pigmentation was analyzed using FIJI. An unpaired t-test was performed for statistical analysis on GraphPad Prism. The p-value is reported for comparison between WT and transgenic caudal fin pigmentation. 19 WT fish and 19 transgenic fish were analyzed. Error bars represent standard deviation.

We next sought to address whether the observed change in the area of pigmentation was due to a change in the average size of the melanophores. We determined that the melanophores in the caudal fins of the Tg fish are smaller than those in the clutchmate WT control fish (Figures 1 and 3), with an average melanophore cell size of 541 µm^2^ for Tg and 746 µm^2^ for WT (Figure 3). Since the Tg fish have increased overall pigmentation, these results imply that there may be a higher number of these smaller melanophores present in the Tg fish. Students could address this experimental question as a next step by quantifying the number of melanophores present in Tg vs WT fish within a certain area of the caudal fin, or by monitoring the proliferation of the melanophores in Tg vs WT via timelapse microscopy. There are certainly multiple avenues of research that could be further pursued in the undergraduate laboratory classroom or during an undergraduate research experience following this two-week laboratory experience.

**Figure 3.**
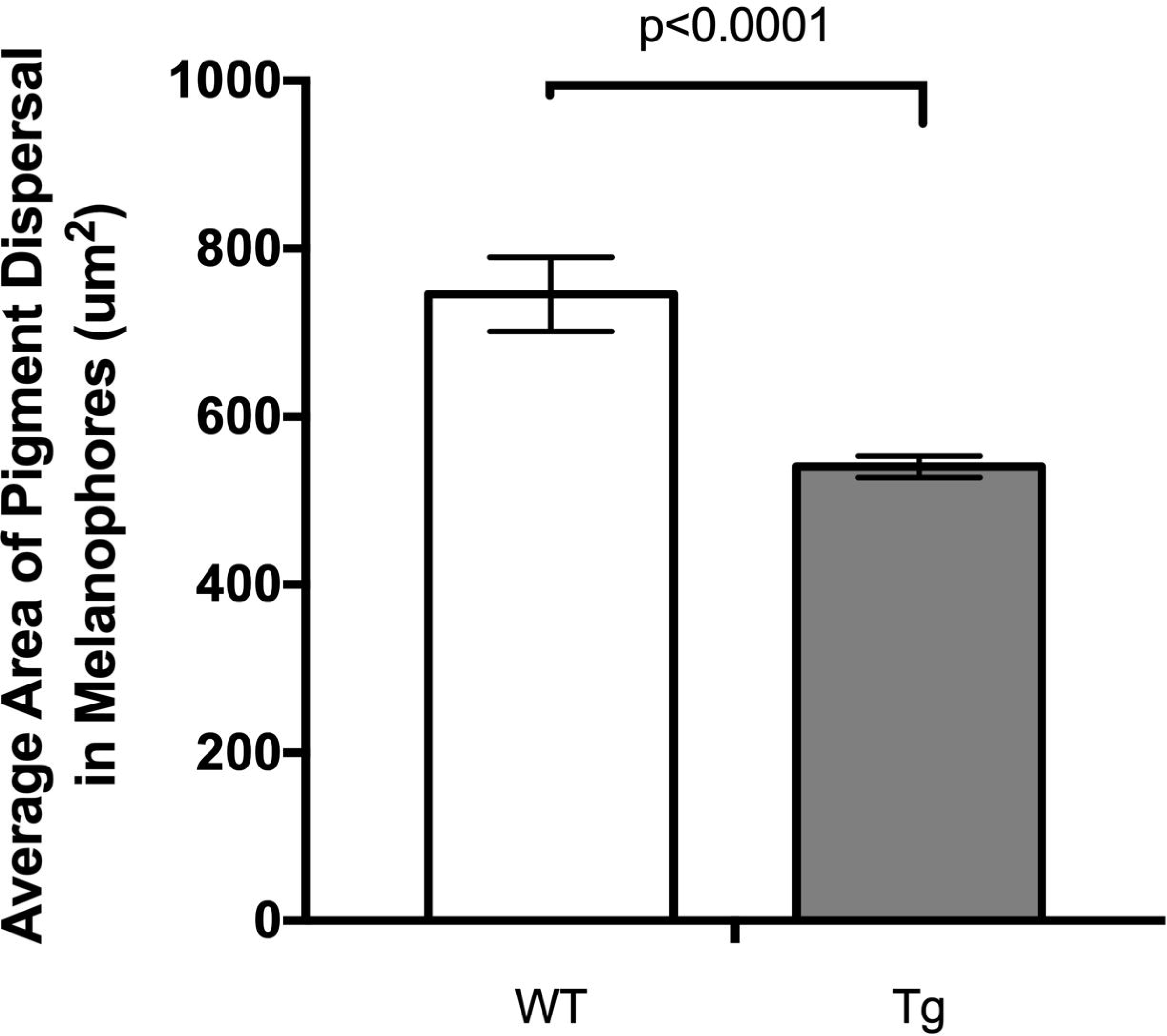
Average area of pigment dispersal (an approximation of cell size) in melanophores in the caudal fins of adult WT and Tg zebrafish. Images of the caudal fins of adult zebrafish were captured on a Nikon SMZ18 stereoscope. Image pigmentation was analyzed using FIJI. An unpaired t-test was performed for statistical analysis on GraphPad Prism. The p-value is reported for comparison between the cell sizes from WT and transgenic caudal fin pigmentation. 204 cells were analyzed from WT fish and 387 cells from transgenic fish. Error bars represent standard error of the mean.

A post-lab survey was administered to all students to assess their confidence levels around the learning objectives of this laboratory (Table 1). Students reported the highest confidence (Likert score 4.63/5) in their ability to use FIJI software to do basic analyses of microscopic images (cell size, area, etc.).

**Table 1.**
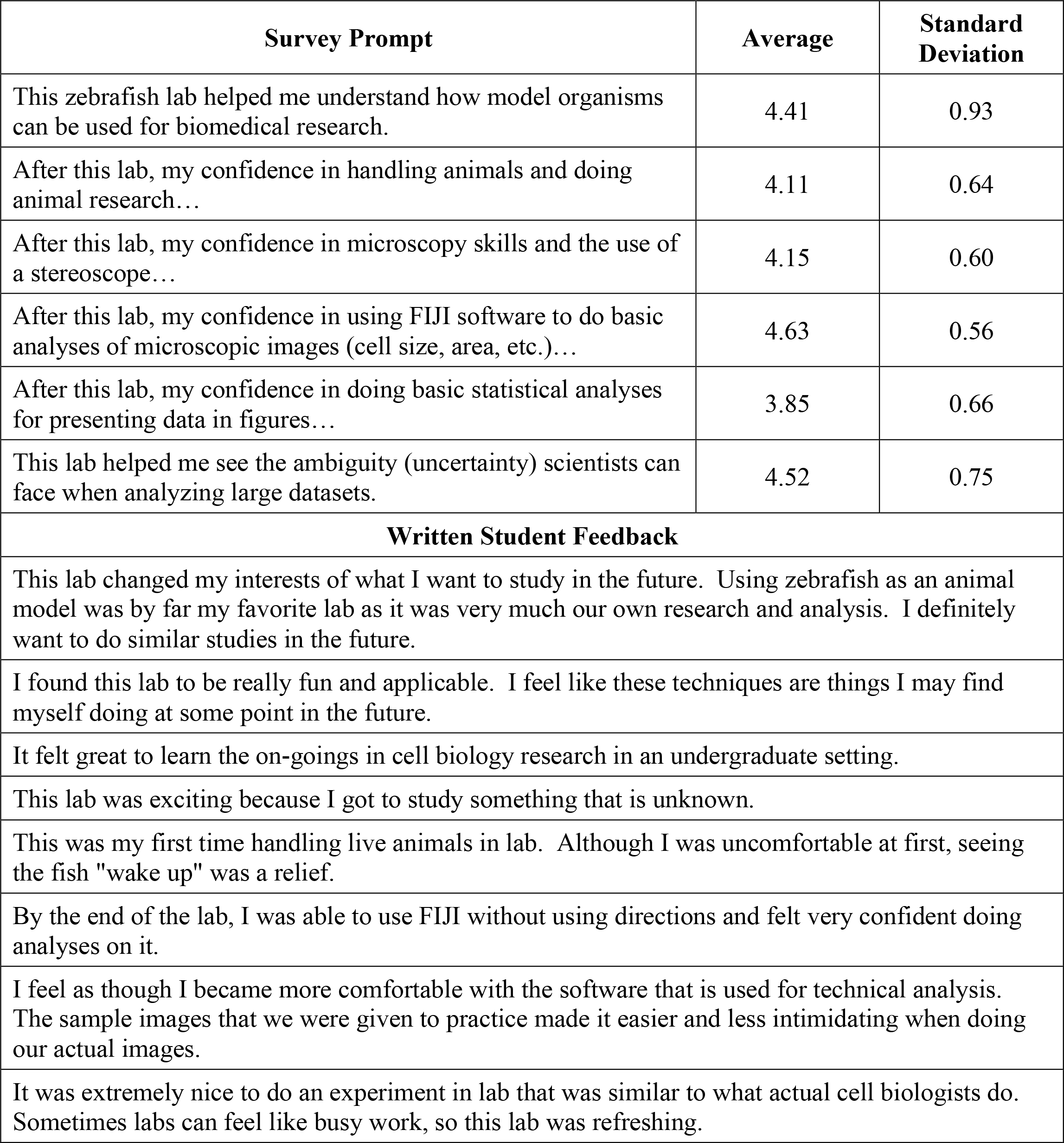
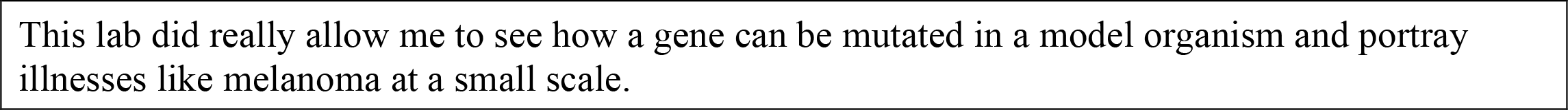
Post-lab survey results of student confidence for each learning objective and written student feedback. Data were collected across two semesters of Cell and Molecular Biology at Carthage College (n=27). Prompts were based on a 5-point Likert scale, where 1=strongly disagree, 2=disagree, 3=neutral, 4=agree, 5=strongly agree, or, 1=(confidence) strongly decreased, 2=decreased, 3=stayed the same, 4=increased, 5=strongly increased. Selected student feedback from the open response prompt within the survey are highlighted.

| Survey Prompt | Average | Standard Deviation |
| --- | --- | --- |
| This zebrafish lab helped me understand how model organisms can be used for biomedical research. | 4.41 | 0.93 |
| After this lab, my confidence in handling animals and doing animal research... | 4.11 | 0.64 |
| After this lab, my confidence in microscopy skills and the use of a stereoscope... | 4.15 | 0.60 |
| After this lab, my confidence in using FIJI software to do basic analyses of microscopic images (cell size, area, etc.)... | 4.63 | 0.56 |
| After this lab, my confidence in doing basic statistical analyses for presenting data in figures... | 3.85 | 0.66 |
| This lab helped me see the ambiguity (uncertainty) scientists can face when analyzing large datasets. | 4.52 | 0.75 |
| **Written Student Feedback** |  |  |
| This lab changed my interests of what I want to study in the future. Using zebrafish as an animal model was by far my favorite lab as it was very much our own research and analysis. I definitely want to do similar studies in the future. |  |  |
| I found this lab to be really fun and applicable. I feel like these techniques are things I may find myself doing at some point in the future. |  |  |
| It felt great to learn the on-goings in cell biology research in an undergraduate setting. |  |  |
| This lab was exciting because I got to study something that is unknown. |  |  |
| This was my first time handling live animals in lab. Although I was uncomfortable at first, seeing the fish "wake up" was a relief. |  |  |
| By the end of the lab, I was able to use FIJI without using directions and felt very confident doing analyses on it. |  |  |
| I feel as though I became more comfortable with the software that is used for technical analysis. The sample images that we were given to practice made it easier and less intimidating when doing our actual images. |  |  |
| It was extremely nice to do an experiment in lab that was similar to what actual cell biologists do. Sometimes labs can feel like busy work, so this lab was refreshing. |  |  |

Importantly, at Carthage College, this class provides biology students with their first encounter with FIJI and any type of biological image analysis. The results of this post-lab survey indicate that students feel quite confident in this type of analysis after this lab, even if this was the first time they encountered this analysis. Their confidence levels may be in part due to the repetitive nature of the image analysis portion of this laboratory. Students are instructed to measure the overall pigmentation in zebrafish at least twice and the calculate cell size measurements for at least 10-15 different cells within each genotype of fish. Practice makes perfect!

Students also agreed (Likert score: 4.52/5) that this lab helped them see the ambiguity (uncertainty) scientists can face when analyzing large datasets. Again, this was one of the first classes that Carthage biology students encountered which asked them to work with and analyze class datasets comprising hundreds of data points. Students sometimes found this experience frustrating, particularly when determining outliers in the large dataset and performing statistical tests. These frustrations were reflected in the post-lab survey too, on the question assessing their confidence in doing basic statistical analyses for presenting data in figures (Likert score 3.85/5). For many of the students, this was their first time performing a t-test in their biology coursework and the first time they were asked to prepare formal scientific figures using Microsoft Excel. Instructors can help ease student anxiety and frustrations by emphasizing that this is a normal part of the scientific process and that professional scientists encounter these difficulties regularly. As a solution to these challenges, instructors can ask their students to discuss their approaches to handling large datasets and statistical tests together as a class, to see if everyone can come to a consensus for analyzing the data using the same consistent methods.

Additionally, this lab helped students feel confident in using microscopes (Likert score 4.15/5), handling research animals (Likert score 4.11/5), and improve their understanding of how model organisms are used in research (Likert score 4.41/5) (Table 1). All of these skills are beneficial for any biology student hoping to attain a research career after graduation. This two-week laboratory module provides students with an introduction to these important skills in the context of an authentic research experience.

The written student feedback (Table 1) from the post-lab surveys was generally positive and emphasized several key features or themes of this laboratory experience:

1.) Future applications of this research to their careers or additional research studies while in their undergraduate education
2.) New skills gained in technical analysis of data and in the use of the software FIJI
3.) Gratitude for doing modern cell biology research and authentic research that has no known answers
4.) Appreciation for zebrafish or other model organisms, and how model organism research can help us better understand biological processes occurring in diseases such as cancer.

While only some of the student feedback has been selected for this manuscript, the general themes and comments have remained consistent across 9 semesters of this laboratory experience and over 250 undergraduate biology students.

In conclusion, this two-week laboratory experience provides undergraduate students with the opportunity to conduct authentic research during a cell biology course. It is in line with the recommended best practices for teaching the scientific process and engaging students in the classroom through active learning^14,27–29^. This laboratory is also readily adaptable to other courses in biology (such as cancer biology or developmental biology) and to other genotypes of zebrafish that an instructor has access to at their institutions. Of importance, particularly for smaller teaching-focused institutions with zebrafish, this lab can be offered with minimal financial investment in terms of necessary reagents or supplies. Students clearly benefit from this research experience, as evident by their increased confidence for conducting research in the laboratory and the attainment of new laboratory and data analysis skills.

## Supporting information

Supplemental Lab Manual

## Acknowledgements

Thank you to the hundreds of students who have taken Cell and Molecular Biology or Advanced Cell Biology at Carthage College over the past 8 years and have provided valuable feedback on this laboratory experience. Your questions and curiosity are inspiring!

