## Supplemental Lab Manual for "Development of an undergraduate cell biology laboratory to assess pigmentation and cell size in a zebrafish model of uveal melanoma"

### Laboratory: Zebrafish Experimentation

#### Investigating the Effects of the GNAQ$^{Q209L}$ Mutation on Melanophores and Pigmentation using Microscopy and FIJI

#### Overview:

In the next two labs you will be conducting authentic research to investigate the effects of the overexpression of a G stimulatory (Gα subunit) protein on pigmented cell morphology in zebrafish. Gα proteins are intracellular proteins that associate with a G-protein coupled receptor. They must exchange GDP for GTP to become active and turn on downstream signaling pathways. Activation of these pathways ultimately leads to effector activation and typically increased cell proliferation and survival. Gα proteins can only become inactive once they hydrolyze GTP to GDP. This inactivation would shut off the signaling pathway.

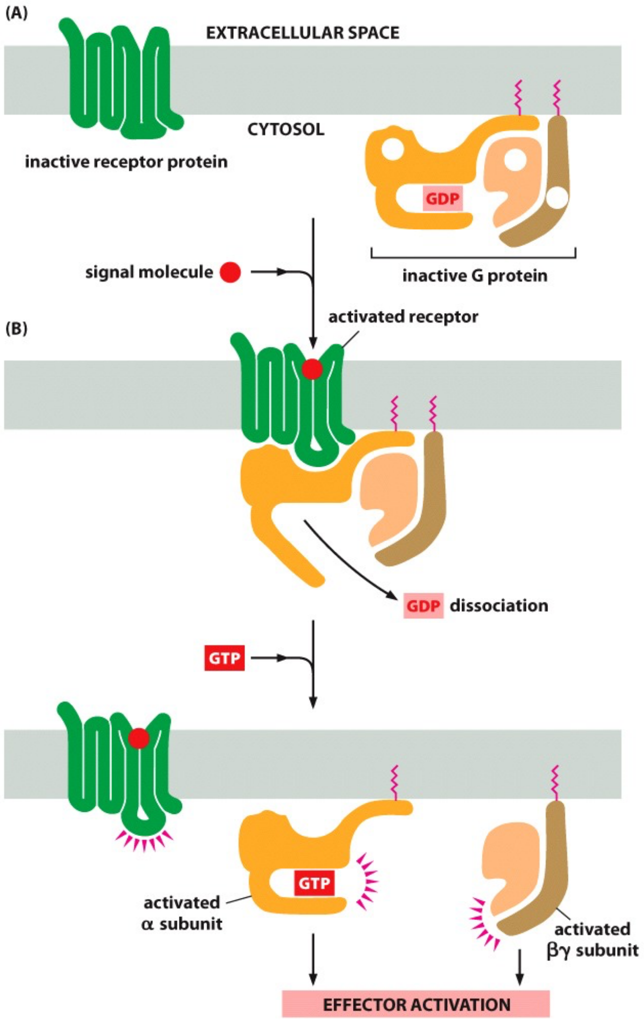


*Image from Essential Cell Biology, 4th Ed., Alberts et al.*

Some of the zebrafish you will be working with in this lab are a transgenic (tg) model for a disease called uveal melanoma. This is a pigmented cancer of the eye or uvea. Melanocytes are the black, pigment producing cells that are affected and become cancerous in this disease. In a transgenic zebrafish model, a foreign gene sequence has been introduced into the genome of the fish. In our particular model, a mutated copy of the human GNAQ gene has been introduced into the zebrafish. GNAQ is one specific type of Gα protein that is mutated in ~83% of humans with uveal melanoma. The mutation most commonly found in the GNAQ gene in humans is a point mutation that changes amino acid 209 from a glutamine (Q) to a leucine (L). A biologist would label this transgenic fish expressing the mutated gene as *Tg(GNAQ$^{Q209L}$)* to indicate amino acid 209 of GNAQ is mutated from Q to L in this transgenic model. Review question: what type of DNA mutation would this particular GNAQ mutation be categorized as (silent, missense, or nonsense)?

The GNAQ$^{Q209L}$ mutation renders this Gα subunit constitutively active. This means it affects the binding site for GTP such that the subunit is unable to hydrolyze GTP to GDP. GNAQ remains bound to GTP constantly when the mutation is present. Therefore, any downstream signaling pathways of the G protein are activated and are turned on in cells with mutant GNAQ. Hypothetically, this is what causes the melanocytes in these uveal melanoma patients to become cancerous and hyper-proliferative.

The main problem is that scientists do not have a good understanding of which specific signaling pathways are activated downstream of GNAQ activation. Without this knowledge, it makes it difficult to understand how the cells become abnormal in the first place, and even more difficult to know how to fix any abnormal signaling pathways or how to inhibit components of the hyperactive pathways with pharmaceutical drugs. Furthermore, there are limited samples available from patients with this disease, necessitating the need for a relevant animal model.

Cancer Cell Biologists have developed a zebrafish model of uveal melanoma with the hope of understanding the signaling mechanisms that cause the cells to become cancerous and the signals that cause the cancer to metastasize to distant sites. Now that the model has been generated, the first step is to characterize the effects of the transgene, mutated GNAQ, on melanophores (pigmented cells) in the zebrafish. This is where you come in. You will research the effects of the transgene on total pigmentation in the zebrafish skin through analysis of pigmented area in an easy to obtain tissue sample, the caudal (tail) fin of the fish. You will also investigate how GNAQ affects individual cells by measuring the area of pigment (melanin) dispersion within each cell.

Given what you know about the role of melanocytes in melanoma, you should formulate a hypothesis for how you think the mutated GNAQ will affect the overall pigment in the zebrafish tail fin and the individual cells themselves. You will compare the analysis of the transgenic fish (genotype *tg:GNAQE32A$^{Q209L}$*) to an age-matched control wild type zebrafish (genotype *Tab 5/14*).

Remember, this is authentic research. This study has not yet been completed. Your instructors do not have the answers, nor do they know what the data should look like. This means it is up to you to carefully collect your data and follow proper experimental procedures. Pay close attention to detail as the data you collect will hopefully be used to advance our knowledge of this disease and could be published in the future.

#### Pre-lab assignment:

https://docs.google.com/document/d/1N6A5-x-EuE41cevXm8t49QX53u3Us9OJ/edit?usp=drive_link&ouid=105380012391681876818&rtpof=true&sd=true

1.) Download and install FIJI (**F**iji **i**s **J**ust **I**mageJ) from the NIH. The software is freely available here: http://fiji.sc/ This is a widely accepted tool used by biologists worldwide for processing of images. If you have a pc, you may have to extract all files before opening the ImageJ application.

2.) Download and save the image found at this link: https://goo.gl/RMb2IA

- Open the installed FIJI software.
- Go to File->Open and click to open the image file you just saved above from google drive.
- Go to Image->Type->8-bit. Type in the pre-lab assignment worksheet (word document) what this selection does to the image (***question 1***).
- Then go to Image->Adjust->Threshold.
- Move the top scale bar to 0 and the bottom scale bar to 50. Type in the same word document what this does to the image (***question 2***).
- Choose Red from the drop-down box on the right side of the threshold menu. What does this do to the image? (***question 3***) Type your answer in your word document.
- Click Set. Then click OK in the Set Threshold Levels box. Then close the Threshold box.
- Go to File->Save As -> Tiff to save the image. Import the image into your word document as part of the assignment (***question 4***). Submit your word document assignment using the submission link for the pre-lab assignment on our learning management system.

3.) Watch the two videos posted on our learning management system.

- The first video (https://youtu.be/8Yg5Ncq_d1w) is a demonstration of the surgical procedure you will perform with the zebrafish in this lab. Watching this video ahead of time greatly improves your success in the lab and may help reduce any fears you have working with animals. The appropriate animal handling techniques will be demonstrated again in lab, but the video allows you to see each step up close, which is often difficult to observe in lab when >10 people are crowded around a small area.
- The second video (https://youtu.be/XJnwbwRVLYA) is an introduction to the parts of the Nikon stereoscope and how to use this dissecting scope to capture digital images of the caudal fin of the zebrafish.

4.) Ensure that you have FIJI properly installed on your laptop and that you will have access to this program during our lab this week and next.

#### Surgical Procedure for Caudal Fin Collection in Zebrafish:

***Note: steps 4 through 9 below must be done quickly and with your full attention. You are working with live animals and you must follow these procedures exactly to minimize any suffering to the animals. Read the directions in their entirety first before starting any procedure. If you have questions, ask before starting.***

1. Put on gloves and spray them with 70% ethanol to disinfect.
2. Ensure that the following supplies are set up at your bench:
    - Paper towel laid across the bench as your clean working surface
    - 70% ethanol spray bottle
    - Kimwipes

- Microscope slides
- Scalpel with retractable safety cover (spray the blade with 70% ethanol to disinfect)
- Slotted spoon (spray with 70% ethanol to disinfect)
- Small net
- Small empty tank for anesthesia
- Tank containing WT or transgenic fish (or a mix of both)

3. Set up a small tank containing anesthetic grade tricaine (0.05% tricaine, 0.0625% sodium bicarbonate, pH 7). Pour just enough anesthesia into the tank so that it is ~¾'' depth. You can prop the tank up on a small level object to help achieve this depth while reducing the amount of hazardous anesthetic solution being used. This solution should be discarded in a hazardous waste container at the end of lab.
4. Identify the fish that you wish to collect first for the experiment. Use a pencil to label the microscope slide appropriately – ie. whether the fish has the wild type phenotype or the mutant ($GNAQ^{Q209L}$ transgenic phenotype). Also label the slide with your name and date. Use a small, clean net to gently scoop the fish out of its tank and into the anesthesia tank. Once in the anesthesia tank, you may have to use your fingers to invert the net if the fish is stuck.
5. Watch the fish carefully for signs of it going under anesthesia. Its breathing should slow down (its gills should contract less over time) and it should rotate to one side or upside down (loss of balance). After a short time (~10 to 15 sec), its muscles should not respond or twitch upon gentle touch with the slotted spoon. Once you notice that the gills are no longer contracting rapidly and have almost slowed down to the point where you can no longer detect movement, remove the fish from the tank using a slotted spoon. It helps to keep the spoon completely horizontal/level when trying to scoop up the fish. Allow any excess anesthetic solution to drip into the tank.
6. Place the fish gently directly on a microscope slide.
7. Use your scalpel to spread out the tail fin (caudal fin) of the fish on the slide.
8. Cut the fin about 1 to 2 mm posterior to the base of the tail (Fig. 1).

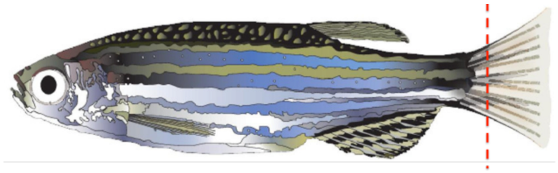

**Fig. 1.** The red dashed line indicates where the caudal fin should be amputated for specimen collection for microscopy. Use a scalpel to cut approximately 1 to 2 mm posterior to the base of the tail; this avoids any cutting of the muscles and vasculature. Zebrafish fins are regenerative and regrow by ~5 days post-amputation.

9. Use the slotted spoon to gently scoop up the fish and return it to its original tank for recovery from anesthesia. It should wake up within a few minutes. You should notice its gills moving and perhaps slight muscle twitches before it starts to swim fully again. If it seems to be taking a while, you can use the slotted spoon to swirl the tank water and increase $O_2$ flow over the gills of the fish. If you are concerned about the recovery of your fish, ask the instructor for guidance.
10. Repeat the above steps for any other fish that you need to collect tail samples from.

#### Image Acquisition using a Stereoscope:

1. Bring your slides to the zebrafish room to collect your microscope images on the Nikon SMZ18 stereoscope. A lab assistant or instructor will be there to instruct you on the proper use of the stereoscope and how to take an image with a scale bar. While you are waiting for your group's turn on the microscope, skip to the image analysis section on the next page.
2. You will collect two images per tail section (Fig. 2): one image of the entire tail at low magnification for calculating total pigmentation, and 1 image of a region of the tail at higher magnification (see red boxes in Fig. 2B) for analysis of pigment dispersion (~ cell size). Make sure to label your files with the appropriate fish background, image number, your name, and date.

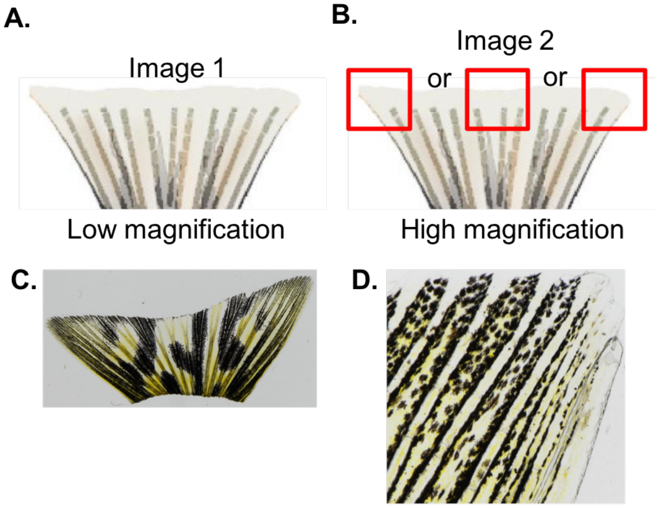


**Fig. 2. Acquisition of caudal fin images on the Nikon SMZ18 stereoscope.** ***A.*** One image of the entire caudal fin will be collected at low (1X) magnification (image #1). ***B.*** One will be collected at the most posterior end of the caudal fin at high magnification. ***C.*** Representative low magnification image for the entire caudal fin (image #1). ***D.*** Representative high magnification image for the posterior right region of the caudal fin (image #2).

### Image Analysis using FIJI:

The first time through these procedures below you should download the sample files (representing an image 1 section and an image 2 section) located at:
[https://drive.google.com/drive/folders/1JY87epsk2RwWRHr69cLLVkXDIqEr3hp1?usp=sharing](https://drive.google.com/drive/folders/1JY87epsk2RwWRHr69cLLVkXDIqEr3hp1?usp=sharing)
Please also download the excel spreadsheet for entering your sample and experimental data for this entire lab. Save the excel file to your computer using the following labeling format: *firstname_lastname_fishgenotype_fish date of birth.* For more details, see step 20 at the very end of this lab manual.

##### *Analysis of total pigmentation in the caudal fin*

1. Open FIJI.
2. Open your tiff image 1 file of the whole caudal fin.
3. Press down the "+" key to zoom in on the image in the region where the scale bar is located. You can use the hand tool 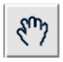 to move the scale bar to the center.
4. Use the straight-line tool 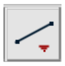 to draw a line across the length of the scale bar.
5. Click Analyze -> Set Scale
6. In the window box that appears, type in um as the units of length. Type in the known distance as whatever your scale was set to (1000 um in this example). Check the global box and then hit ok to set the scale for the image. This allows FIJI to convert the pixels in the image to a size in microns (um), which will be useful when we calculate area in $um^2$. Click OK. (image on next page)

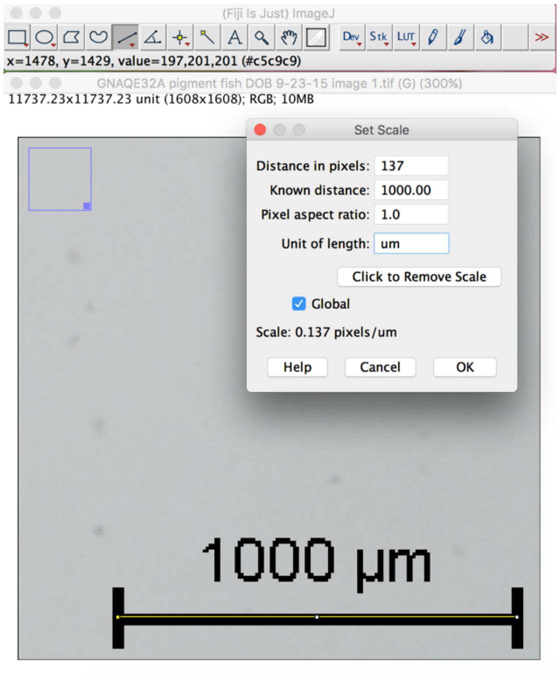


7. Click Image -> Type -> 8-bit to change the image to B&W for the analysis. This makes calculations of area and intensity simpler because the computer is measuring just one color (black). You can use the "-" key to zoom back out to see the entire caudal fin in the window.

8. Use the freehand selection tool 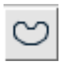 and your mouse to trace around the outside of the tail. Keep the mouse button held down as you are tracing. Be precise, this takes patience and a steady hand. It should look something like the image below when done. The tail is outlined by your yellow freehand selection.

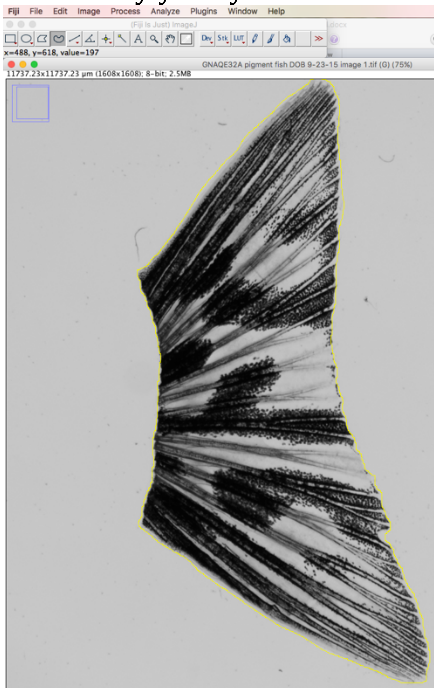


9. Press the "x" key to cut the selection. The region of tail you selected should disappear and be replaced with a black area.

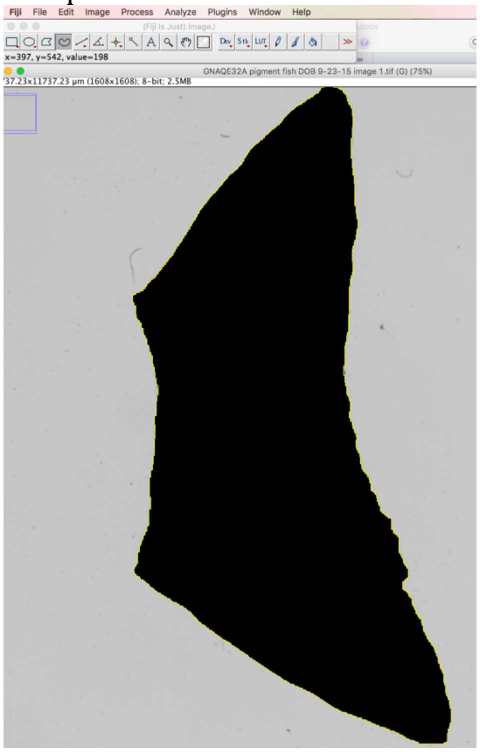

10. Go to File-> New -> Image…
11. In the new image window, type in the image size of the original file you had opened (it should be 1608 pixels x 1608 pixels if you used the Nikon SMZ18). Make sure 8-bit is selected as the type. Make sure White is selected as the background under the Fill with: drop down box. Click OK.

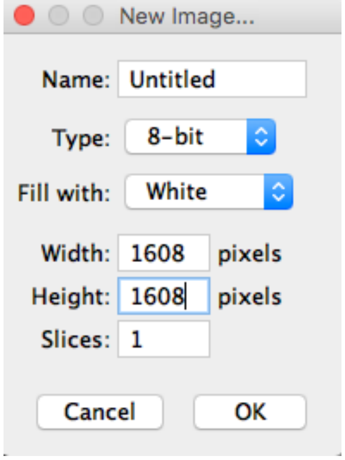


12. Press the "v" key to paste the selection in the new image window.
13. With the tail image still selected by the yellow outline, you should next click Analyze -> Set Measurements. If you lose the selection at any point you can go to Edit –> Selection -> Restore Selection.
14. In the Set Measurements window, check the boxes next to Area, Area fraction, and Limit to threshold. Ensure that Decimal places is set to 3. Click OK. (image on next page)

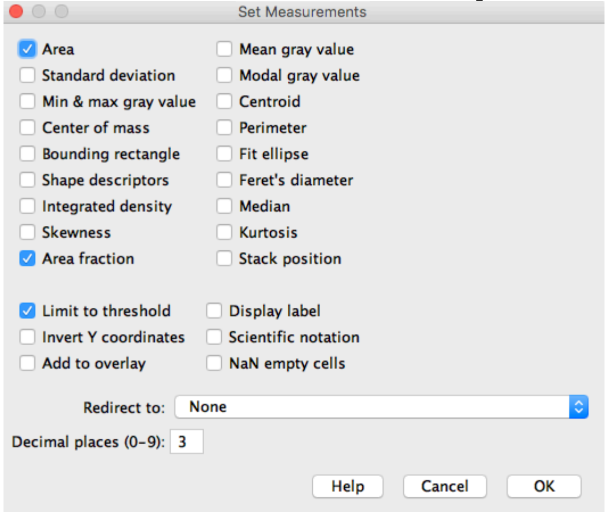


15. Then go to Analyze –> Measure, to calculate the **total area of the selected tail in um²**. Copy and paste this row of measurements from the results window into the excel data file.

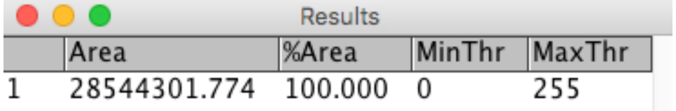


16. Now click Image -> Adjust -> Threshold.
17. Select Red from the drop down box on the right side of the Threshold window. (It might be displaying B&W as the default. Change it to Red.)
18. Open your original tiff file in whatever program your computer has for viewing images and keep it to the right of your FIJI file as a reference image to help you determine if you have oversaturated your selection of pigment in the tail.

19. Now move the bottom Threshold slider to select the majority of the black pigmented area in the FIJI caudal fin image as red. Compare your selection to the original fin image in the tiff file. You don't want to oversaturate your selection and select regions that were not initially black or pigmented. You also don't want to under-select the pigmented region and not have it be accounted for in the calculation for pigmented area. In this particular example below, a threshold setting of 66 seems to be just right for selecting the majority of the black pigmented regions in the tail while ignoring the yellow pigmented regions and the unpigmented bony ray structures (bones) in the fin.

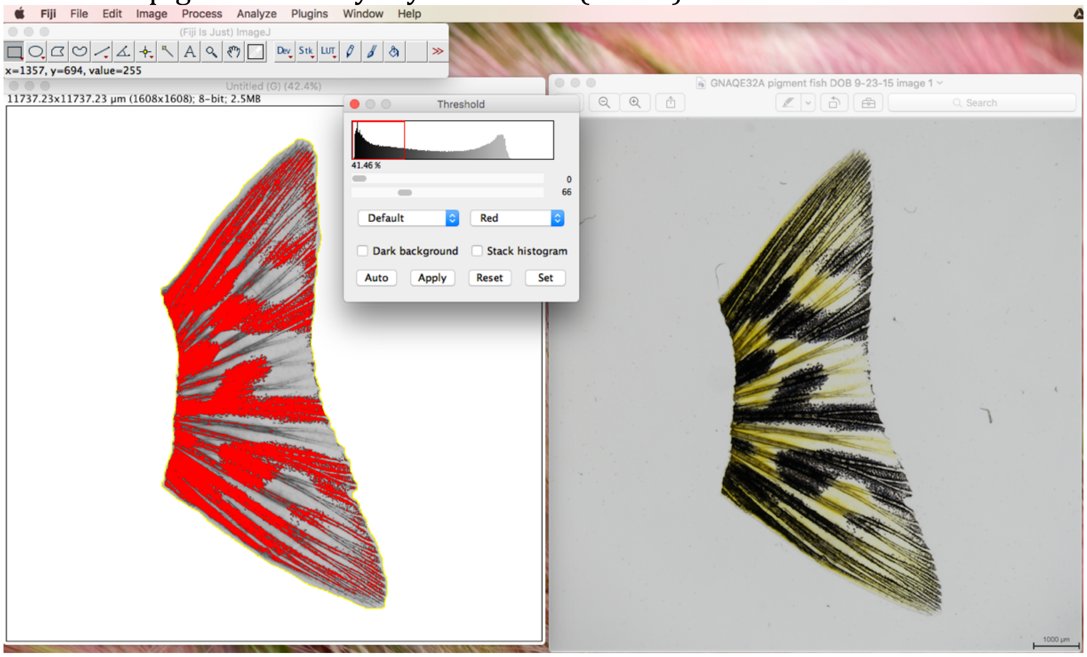


20. Click Set. A window will appear that says Set Threshold Levels. Click OK.

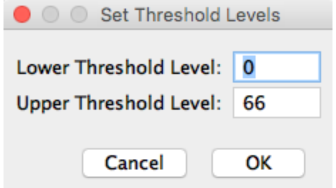


21. Now click Analyze -> Measure, in order to calculate the total area of the thresholded/red part of the image. This represents the **total area of pigmentation in um² in the caudal fin**. The results will appear in the Results window as the next row.

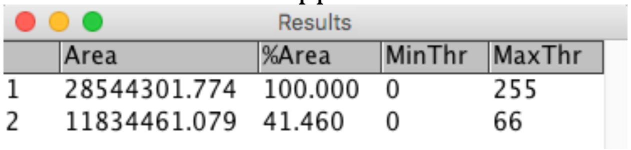
Results

| | Area | %Area | MinThr | MaxThr |
|---|---|---|---|---|
| 1 | 28544301.774 | 100.000 | 0 | 255 |
| 2 | 11834461.079 | 41.460 | 0 | 66 |

22. Copy and paste this row of values into your spreadsheet. In your spreadsheet, manually calculate the percent red area (results #2 area measurement) out of the total tail area (results #1 measurement). You should get roughly the same value as what FIJI displayed as %area in the most recent measurement for the thresholded red selection. If you don't get the same value, double-check your procedure again. Ask an instructor for assistance if you cannot figure out what went wrong.
23. You have now calculated the area of pigmentation in the tail in um² and the percent % of the tail that is pigmented. Save your spreadsheet.

*Analysis of pigment dispersion (~cell size) in the caudal fin*

To start, think about why we consider this a measurement of the area of pigment dispersion within a cell and not a measurement of cell size. What would you need to measure or be able to detect to calculate cell size or the area of a cell?

1. Open FIJI.
2. Open your tiff image 2 file of a magnified region of the tail fin.
3. Press down the "+" key to zoom in on the image in the region where the scale bar is located. You can use the hand tool to move the image around to that image.
4. Use the straight-line tool 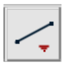 to draw a line across the length of the scale bar.
5. Click Analyze -> Set Scale
6. In the window box that appears, type in um as the units of length. Type in the known distance as whatever your scale was set to (100 um in this example). Check the global box and then hit ok to set the scale for the image. This allows FIJI to convert the pixels in the image to a size in microns (um), which will be useful when we calculate area of pigment dispersion (cell size) in $um^2$. Click OK.

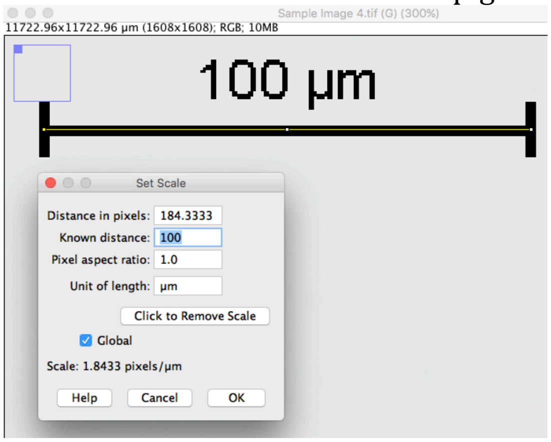


7. Click Image -> Type -> 8-bit to change the image to B&W for the analysis. You can use the "-" key to zoom back out to see the entire caudal fin in the image.
8. Open your original tiff file for image 2 in whatever program your computer has for viewing images. Keep this image file to the right of your FIJI file as a reference image to help you determine if you have oversaturated your selection of pigment dispersion in the tail.
9. Click Image -> Adjust -> Threshold.
10. Make sure Red is selected in the drop-down box. Then use bottom slider bar to adjust the saturation so that the red color selects the majority of the black, pigmented cells. In the example below, a threshold of 92 seems to be just right to select the majority of black, pigmented cells seen in the reference image. (image on next page)

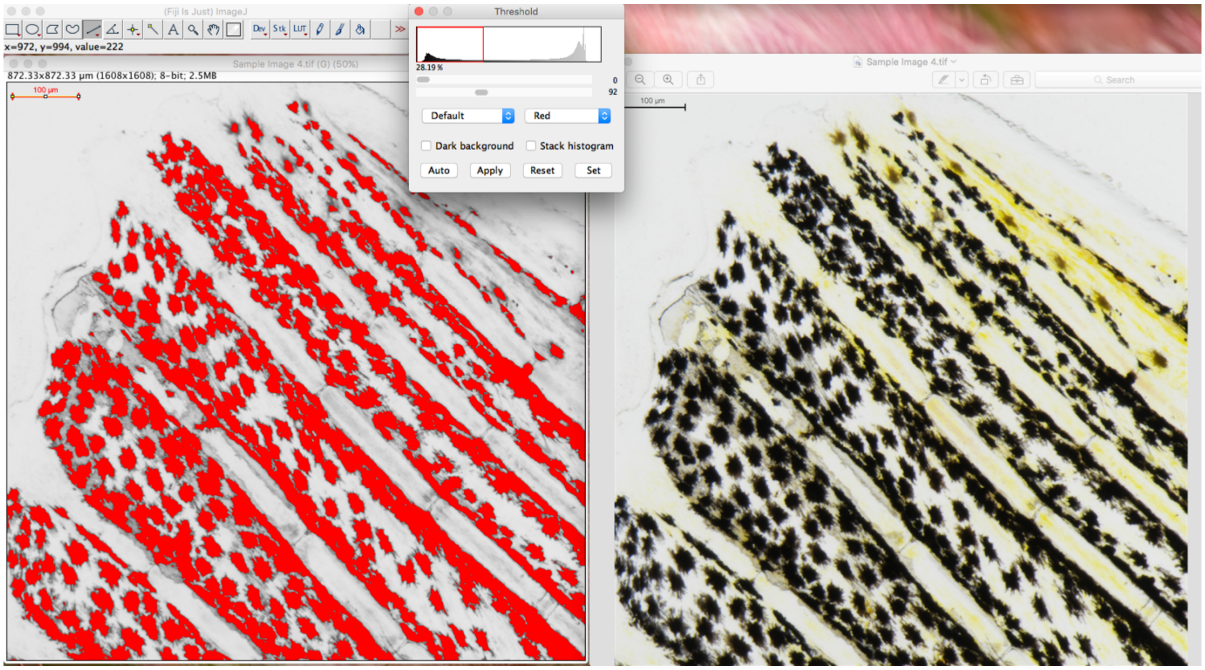


11. Click Set in the Threshold window.  Confirm that your bottom slider value is displayed as the Upper Threshold Level in the Set Threshold Levels window.  Then click OK.

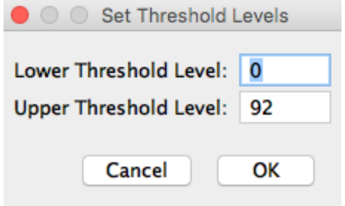


12. Next click Analyze -> Set measurements.
13. Check the boxes next to Area and Limit to threshold.  Ensure decimal places is set to 3. Click OK.

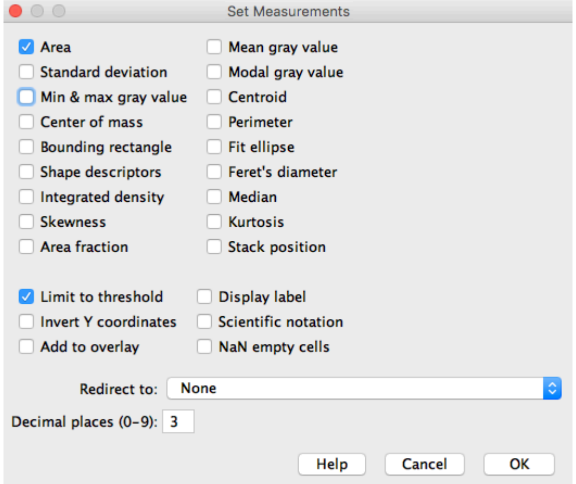


14. Now you can analyze the area of each cell (pigment dispersion).  Use the freehand selection tool 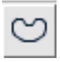 and hold our mouse button down to draw around the outside of each cell.  Try your best not to get the neighboring cells in your selection, as it would change your area calculation.  If the cells are clumped together, do not calculate their area.  Only select cells for which you can see a clear boundary representing a single cell.

15. Once you have the entire area selected around the cell, go to Analyze -> Measure to bring up the results. Copy and paste the results line into your excel file. You have now calculated the area pigment dispersion within a single cell.
16. With your cell still selected in Fiji, then click on Analyze -> Tools -> ROI Manager. Click Add or hit the "t" key. This will save your current freehand selection of the cell so we can return to it later and look at all of the cells we analyzed in a given image. We can re-analyze the selections later if we need to too. Check the boxes next to Show All and Labels. This will keep your selections visible in your image and will assign a number to each selection so you know the order in which you measured them. Record the ROI Manager # for each freehand selection in the appropriate column in the excel spreadsheet too.
17. Repeat the above steps for at least 15 randomly selected cells in the image. If time allows, try to analyze all of the cells in the image for which you can see clear boundaries (ie. cells for which there is white space surrounding all sides of the red selection). Be patient, this will take some time, which is why we are spending two weeks on this lab. This data may be used in a future publication, so consistent analysis and attention to detail are key.

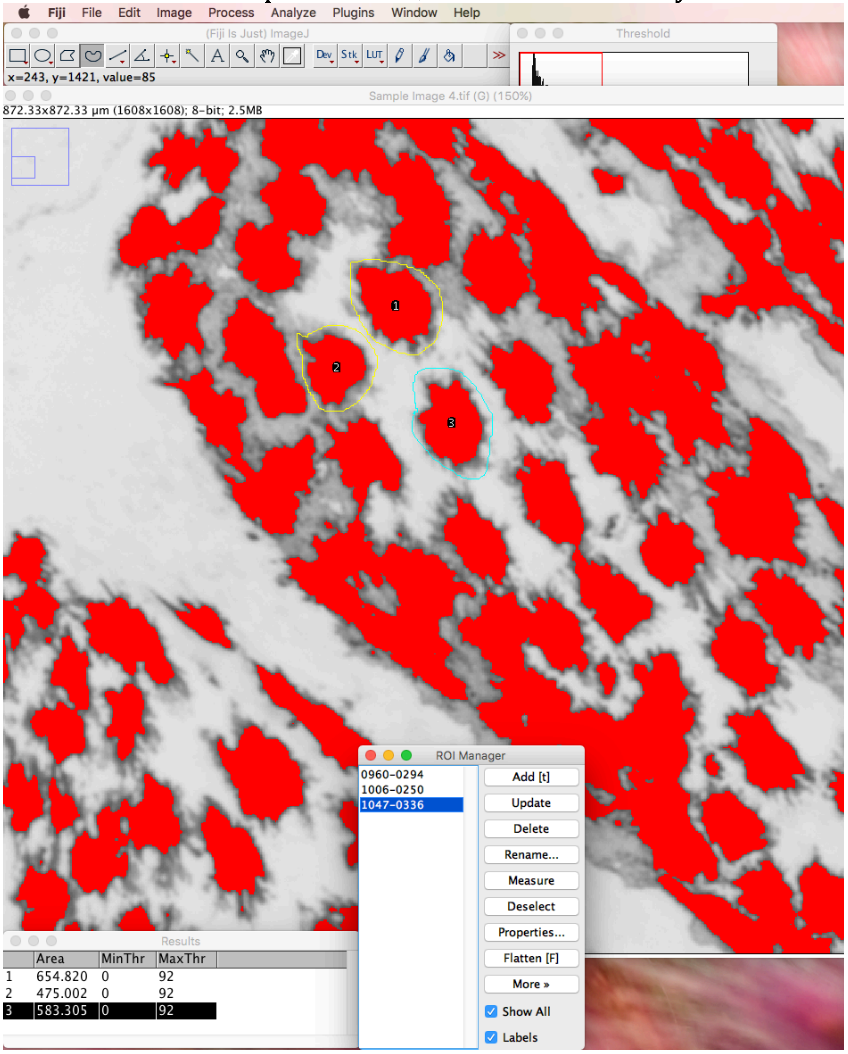


18. Once you have analyzed all of the cells in your first image, save your image file (with the scale and thresholds set) as a new .tiff file by clicking File->Save As -> Tiff.... Label the file in the following format: *firstname_lastname_fishgenotype_image2_fishDOB*, where fish genotype refers to either WT or GNAQE32B or GNAQE32A fish (whichever fish you chose to amputate).

19. Before closing out of FIJI, save your ROI files. In the ROI manager window, select all of the file numbers at once using the shift key. Then click the "more" box and save. Save these files as the same descriptive name, but indicate they are ROI files at the end of the file name as such: *firstname_lastname_fishgenotype_image2_ROIs* They will save as a .zip file.
20. Save your excel spreadsheet file in the following format: *firstname_lastname_fishgenotype_fish date of birth.* The date of birth should be written MMDDYY, so 110619, for instance.
21. At the end of the day, make sure to upload your original stereoscope image files, your FIJI image files, the ROI zip files, and excel spreadsheet to a shared class folder so the instructor has access to all of your data for next week's analysis during lab. **<u>This should be completed by 8AM two days before our next lab.</u>** Make sure each uploaded file is labeled appropriately as instructed throughout this lab manual. (Include a link to a shared google drive folder that contains subfolders for each type of file).
